# Fluorescent Nanosensors Reveal Dynamic pH Gradients During Biofilm Formation

**DOI:** 10.1101/2020.07.31.230474

**Authors:** Birte Blunk, Mark Perkins, Veeren M. Chauhan, Jonathan W. Aylott, Kim R. Hardie

## Abstract

Understanding the dynamic environmental microniches of biofilms will permit us to detect, manage and exploit these communities. The components and architecture of biofilms have been interrogated in depth, however, little is known about the environmental microniches present. This is primarily because of the absence of tools with the required measurement sensitivity and resolution to detect these changes. We describe the application of ratiometric fluorescent pH-sensitive nanosensors, as a novel tool, to observe physiological pH changes in biofilms in real-time. Nanosensors comprised two pH-sensitive fluorophores covalently encapsulated with a reference pH-insensitive fluorophore in an inert polyacrylamide nanoparticle matrix. The nanosensors were used to analyse the real-time three-dimensional pH variation for two model biofilm formers: (i) opportunistic pathogen *Pseudomonas aeruginosa*, and (ii) oral pathogen *Streptococcus mutans*. The detection of sugar metabolism in real time by nanosensors provides a potential application to identify novel therapeutic solutions to improve oral health.

## Introduction

The biofilm communities that bacteria form on surfaces are dynamic and complex. The architecture of biofilms and the identity of extracellular components have been characterised *in vitro* for a variety of biofilms, although there is discussion in the literature about how representative this is of *in vivo* bacterial communities ^1,2,3,4,5^. Whilst there is evidence for fluid channels and microcolonies within *in vitro* biofilms, the characterization of the environmental microniches these create has been limited due to the absence of analytical tools that are suitably sensitive at the required resolution. In this article we demonstrate the application of fluorescent nanosensors capable of dynamic pH monitoring of the environmental microniches within biofilms using two important pathogens.

Firstly, we selected the Gram-negative bacterium *Pseudomonas aeruginosa*, an opportunistic pathogen, causing infections in immunocompromised patients including those suffering from burns, wounds and Cystic Fibrosis (CF) ^6,7,8,9^. These infections are challenging to treat, due to the intrinsic antibiotic resistance of *P. aeruginosa*, which contributes to *P. aeruginosa* infections being the leading cause of morbidity for CF sufferers. *P. aeruginosa* is prevalent in the environment creating numerous potential reservoirs of infection. *P. aeruginosa* exerts its pathogenicity through the production of several factors including those shown to play a key role in biofilm attachment such as extracellular DNA ^10,11,12,13^. Biofilm formation by *P. aeruginosa* is particularly important for it to establish chronic infections and has therefore been studied intensively.

Secondly, we chose the Gram-positive oral pathogen *Streptococcus mutans.* Oral biofilms play a crucial role in the aetiology of oral diseases, such as dental caries and periodontitis, which can lead to increased economic burden and reduced quality of life ^14^. Oral biofilms are characterised by a bacterial shift from early colonisers of the tooth towards increasingly acid-producing (acidogenic) and acid tolerant (aciduric) species ^15^. *S. mutans* is both an aciduric and acidogenic bacterium and a pre-dominant species in late stage oral biofilms ^16,17^. The interest in oral bacteria such as *S. mutans* lies in their participation in dental caries, the most common infection affecting humans, through the production of organic acids ^18^. Initially, *S. mutans* uses sucrose as a substrate to synthesise glucan *via* glucosyltransferases, in order to provide anchoring sites on the tooth’s enamel. This enables bacteria to continue to colonise and establish more biofilm ^19,20,21^. Once established, *S. mutans* ferments available carbohydrates, readily found in a sugar-laden diet, which results in the production of organic acids ^16^. This acidification leads to the formation of dental caries ^22,23,24^. Robust, well characterised biofilm models exist for both *P. aeruginosa* and *S. mutans*, making them an ideal choice for this study.

An environmental characteristic unlikely to be static and homogeneous in biofilms is pH, making it an attractive option to quantify spatially in real-time. It is also important to characterize the environmental pH of microorganisms because it can influence many different physiochemical properties such as polymer-polymer, ion-polymer and macromolecule-polymer interactions ^25^. Furthermore, the formation of chemical gradients (such as in pH, redox potential and ions) within the biofilm community can provide indications about microbial metabolism ^26,27,28,29,30^.

In the literature, the observations of the relationship between pH and biofilm formation differ. For example, Ahmed *et al.* demonstrated that a trigger of biofilm formation in some bacteria, autoinducer-2-signalling (AI-2), is temperature dependent in *Streptococcus intermedius* but not pH dependent ^31,32,33^. In contrast, Sun *et al.* showed that a low pH together with glucose could improve AI-2 activity and that the addition of a low concentration of boracic acid induces the synthesis of AI-2 ^34^. Although this could be relevant for *S. mutans*, *P. aeruginosa* does not synthesize AI-2. However, like many other bacteria, some of the molecules which *P. aeruginosa* produces, such as signal molecules for quorum sensing (QS) to mediate population density dependent cell-cell communication, are *N*-acylhomoserine lactones (AHLs). AHLs have been implicated in biofilm formation for many bacterial species. For instance, Yates *et al.* showed that the lactonolysis of AHLs is pH dependent, with ring closure only taking place at reduced pH levels ^35^, whereas Hostacka *et al.* demonstrated an accelerated biofilm production for *P. aeruginosa* at pH 7.5 and pH 8.5 in comparison to pH 5.5 using crystal violet quantification ^36^. A similar behaviour was described by Stoodley *et al.* who observed that *P. aeruginosa* biofilm thickness fell to 68% at a low pH of 3 ^37^.

In contrast, studying the Gram-positive bacteria *Staphylococus aureus* and *Staphylococus epidermidis*, Nostro *et al.* found biofilms were 2.5 to 3 times thinner at high pH levels suggesting this to be a possible way to control biofilm formation in this species ^38^. Studies of *Streptococcus agalactiae*, whose infection can result in meningitis, have shown an accelerated biofilm formation in highly acidic conditions ^39^. For other group B streptococcal isolates, the same behaviour was found in vaginal infections of pregnant women where clinical isolates showed a higher biofilm formation rate at a vaginal pH of 4.5 in comparison to pH 7 ^40^.

There are a variety of methods to measure intracellular pH, including nuclear magnetic resonance spectroscopy (NMR), microelectrodes, surface-enhanced Raman scattering (SERS) and two-photon excitation microscopy (TPE) ^28,41,42,43^. However, those technologies are not accessible to all laboratories and the resolution of some of the methods is not yet comparable to fluorescent-based measurements ^25,43,44^. Therefore, nanosensors containing commercially available fluorophores have aroused increasing interest and are most widely implemented in approaches to pH measurement.

Nanosensors are nanoparticles that can be used for the detection and measurement of small environmental changes. Fluorescent nanosensors consist of spherical polymer particles carrying fluorophores (signal transducer) generating fast, bright responses that can be measured using real time fluorescent microscopy ^25,45^. Incorporation of different pH-sensitive fluorophores in the correct ratio in a single nanosensor can enable the detection of the full physiological pH spectrum from pH 3.5 to pH 7.5 ^46,47,48^.

A range of different pH nanosensors has already been applied in biological research, for example in cancer research where it has been suggested that low oxygen concentration and low pH increase the potential of cancer metastasis and reduce the treatment prognosis for patients with advanced stage cancers ^49^. Within microbiology, silica based nanosensors were used to analyse pH microenvironments in microbial biofilms of *Escherichia coli* ^44^. Hidalgo *et al.* could show local variation in pH from the neutral outside surface of the film to the acidic core. Nanosensors with a single fluorescent ratiometric pH-sensitive probe were used to reveal pH heterogeneity throughout a *P. aeruginosa* biofilm ^25^. A slightly acidic biofilm environment was reported with a pH range between 5.6 within the biofilm and 7.0 in the bulk fluid ^25^. More recently, Fulaz *et al.* reported the use of pH-sensitive nanosensors to detect pH gradients in biofilms, but did not extend their study to real time analysis of the environmental microniches under flow conditions ^50^.

As different laboratories use a variety of nanoparticles, characterisation of the interaction of nanoparticles with bacteria and biofilm components is required. For their use in biological systems, it is important that the nanoparticles do not alter the metabolism and behaviour of the cells.

The development of nanoparticles has also been mentioned as a prevention method for application in early stages of diseases ^51^. Biodegradable polymer based nano- and microparticles can, for example, be used to encapsulate antibiotics to ensure a targeted release of the antimicrobial substance ^52,53,54^. Therefore, it is important to understand the interaction between the nanoparticles and the biofilm and to establish the penetration of nanoparticles into the biofilm. Penetration usually occurs by diffusion where the nanoparticles are interacting with either the bacteria or the biofilm matrix components ^52,55,56^. The diffusion characteristics of the nanoparticles mostly depend on their size and charge but are also influenced by pore size of the biofilm, the charge and chemical gradient of the biofilm matrix, and the hydrophobicity of the environment ^52,56,57,58^. An increasing size and negative charge of carboxylated silver nanoparticles has been reported to reduce their diffusion into *Pseudomonas fluorescens* biofilms ^56^. A similar behaviour was seen for anionic liposomes added to a *Staphylococcus aureus* biofilm, where the liposomes could not penetrate the biofilm, possibly due to the negative surface charge of the bacteria ^59^.

From the repertoire of nanosensors available to us, we selected polyacrylamide-based nanoparticles due to their small size. This differed from the nanosensors used by Fulaz et al., which were silica based ^50^. To enable a physiological range between pH 3 and pH 8 to be measured and visualised, two pH-sensitive fluorophores, Oregon green (OG) and 5(6)-carboxyfluorescein (FAM), were used. These fluorophores together with a reference pH-insensitive fluorophore, 5(6)-carboxytetramethylrhodamine (TAMRA), were covalently linked to a polyacrylamide nanoparticle matrix to synthesise ratiometric pH-sensitive nanosensors. To evaluate the influence of charge, both neutral and positively charged versions of the nanosensors were produced.

The fluorescent pH-sensitive polyacrylamide-based nanoparticles were used to investigate the pH microenvironments of *P. aeruginosa* biofilms. The penetration and interactions of the nanosensors within the biofilm were characterized, and the use of nutrient flow and time-lapse imaging enabled real-time reporting of pH modulation within a biofilm to be monitored. We reveal an acidic core within microcolonies and acidification in the downstream flow. In addition, this study highlighted the advantages of nanosensors by using a biologically relevant model for oral health; detecting the effects of various carbohydrates on the external pH of a culture and providing the potential application of this methodology in the development of novel treatments for dental caries.

## Results

From the broad selection of nanoparticles available to us ^60,61^, polyacrylamide nanoparticles were selected. The basis for this choice was to have nanosensors available that were small enough to penetrate deep into biofilms and report with the sensitivity required for high resolution imaging as well as the option to alter surface characteristics and thereby manipulate interactions with the extracellular matrix. Polyacrylamide nanoparticles can be prepared at a smaller size (34.77 +/− 4.63 nm, see Table 1) to those based on silica that were used previously (47 +/− 5 nm) ^50^, and with different surface charges. From our two model bacteria, *P. aeruginosa* was chosen for our initial optimization experiments due to the availability of both static and flow biofilm models.

**Table 1.** Characteristics of the nanosensors synthesized in this study. The size, polydispersity index (PDI) and zeta potential values of positively charged polyacrylamide nanosensors, positively charged polyacrylamide nanosensors without dyes and neutral polyacrylamide nanosensors, determined by a MalvernTM DLS instrument.

|  | Positively charged<br>polyacrylamide<br>nanosensors (n=3) | Positively charged<br>polyacrylamide<br>nanoparticles without<br>dyes (n=1) | Neutral polyacrylamide<br>nanosensors (n=1) |
| --- | --- | --- | --- |
| Size (nm) | 34.77 +/- 4.63 | 46.41 | 37.51 |
| PDI | 0.168 +/- 0.036 | 0.167 | 0.096 |
| Z-potential (mV) | 15.6 +/- 3.85 | 17.7 | -4.75 |

*PDI* polydispersity index, *n* number of preparations analysed

### The addition of positively charged pH-sensitive polyacrylamide nanosensors to PAO1-N WT results in a thicker and more robust biofilm

Having settled on *P. aeruginosa* as our model system for initial optimisation experiments, strain PAO1-Nottingham (PAO1-N) was initially chosen to analyse the influence of nanosensors on biofilm growth. The nanosensors were suspended in the cell culture prior to the incubation with bacteria. The resultant biofilms, grown with and without nanosensors under static conditions, were analysed after 48 h using fluorescence Confocal Laser Scanning Microscopy (CLSM, Fig. 1a). The neutral pH-sensitive nanosensors dispersed throughout the *P. aeruginosa* biofilm without altering their architecture or thickness (Fig. 1a). Positively charged pH-sensitive nanosensors containing acrylamidopropyltrimethylammonium chloride (ACTA) were exchanged with the neutral pH-sensitive polyacrylamide nanosensors. Interestingly, biofilms grown together with the positively charged pH-sensitive polyacrylamide nanosensors were approximately four times thicker (40 μm) compared to biofilms grown in the absence of nanosensors or with neutral pH-sensitive nanosensors (~10 μm) (Fig. 1a). The general structure of the biofilms grown with positively charged pH-sensitive nanosensors was also notably different, showing a very dense structure. To exclude the possibility that the dyes incorporated within the polymer matrix of the positively charged pH-sensitive nanosensors were responsible for the enhanced biofilm formation, positively charged polyacrylamide nanoparticles featuring the same characteristics in size and charge were produced without inclusion of the fluorophores (Table 1). The addition of these nanoparticles led to the similar increases in biofilm thickness and robustness as the positively charged nanosensors, suggesting that the charge of the particles was responsible for the change in the biofilm formation.

**Fig. 1.**
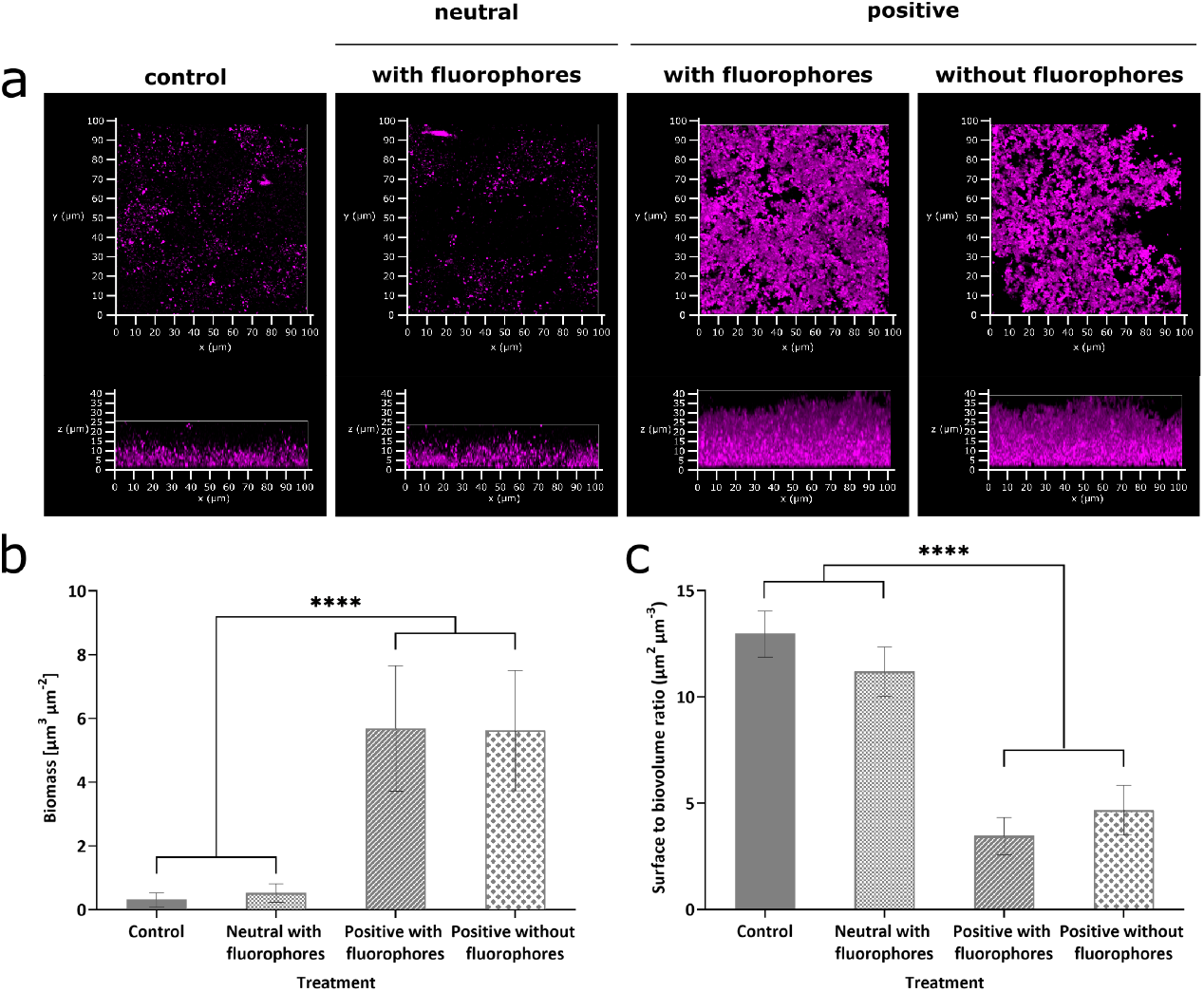
The charge of nanosensors can influence the biofilm formation of *P. aeruginosa* PAO1-N. Differently charged nanoparticles at 1 mg mL^−1^ were incubated with *P. aeruginosa* PAO1-N under static conditions. Quantitative Comstat analysis of the biofilm images were performed using ImageJ. (**a**) Representative confocal images of *P. aeruginosa* PAO1-N stained with CellMask™ without the addition of nanoparticles (left panel), grown with neutral polyacrylamide nanosensors (left central panel), with positively charged polyacrylamide nanosensors containing fluorophores (right central panel) or with positively charged polyacrylamide nanoparticles without fluorophores (right panel). The top row is a 3D view from the top of the biofilm, the bottom row shows a 3D view from the front of the biofilm. Graphs represent (**b**) biomass and (**c**) surface to biovolume ratio. Error bars represent standard deviation measured for different biofilm images, where n=10 (control), n=7 (neutral nanosensors and positive nanoparticles without dyes) and n=8 (positive nanosensors).

Quantitative analysis of the images using Comstat, an open source programme used to analyse image stacks of biofilms ^62,63,64^, revealed that there is no significant difference between the biofilms produced in the absence of nanoparticles or with neutral nanosensors (Fig. 1b and c). Furthermore, a significantly lower biomass for biofilms grown without nanosensors or with neutral nanosensors was confirmed (p < 0.0001, Fig. 1b). In contrast, no significant difference in biofilm biomass was detected between formation in the presence of positively charged pH nanosensors and the positively charged nanoparticles without dyes. In addition, a significantly higher surface area to biovolume ratio for biofilms grown without nanosensors or with neutral nanosensors than those grown with positively charged nanosensors was observed (p < 0.0001, Fig. 1c).

### The concentration of the nanosensors is critical to achieve thicker, more robust biofilms

The effects of nanosensors upon *P. aeruginosa* were assessed by introducing different concentrations of nanosensors to planktonic cell cultures. To detect the effects, the growth of the bacteria was monitored via the OD_600_ every 15 min for 24 h in 96 well TECAN plates. No growth inhibition was observed for nanosensor concentrations below 25 mg mL^−1^ (Supplementary Fig. 1). To investigate whether higher concentrations of nanosensors would lead to an even thicker biofilm than using the normal previously selected concentration of 1 mg ml^−1^, biofilms were grown under the same conditions using 10 mg mL^−1^ or 20 mg mL^−1^ positively charged nanosensors. CLSM revealed that a higher concentration of positively charged nanosensors does not seem to further enhance the biofilm formation, instead the thickness is reduced compared to when 1 mg mL^−1^ nanosensors were introduced (Fig. 2). This was again underlined using Comstat for quantitative analysis of the different conditions, where a significant reduction in biomass was observed for biofilms grown with 10 or 20 mg mL^−1^ compared to 1 mg mL^−1^ nanosensors (p < 0.0001).

**Fig. 2.**
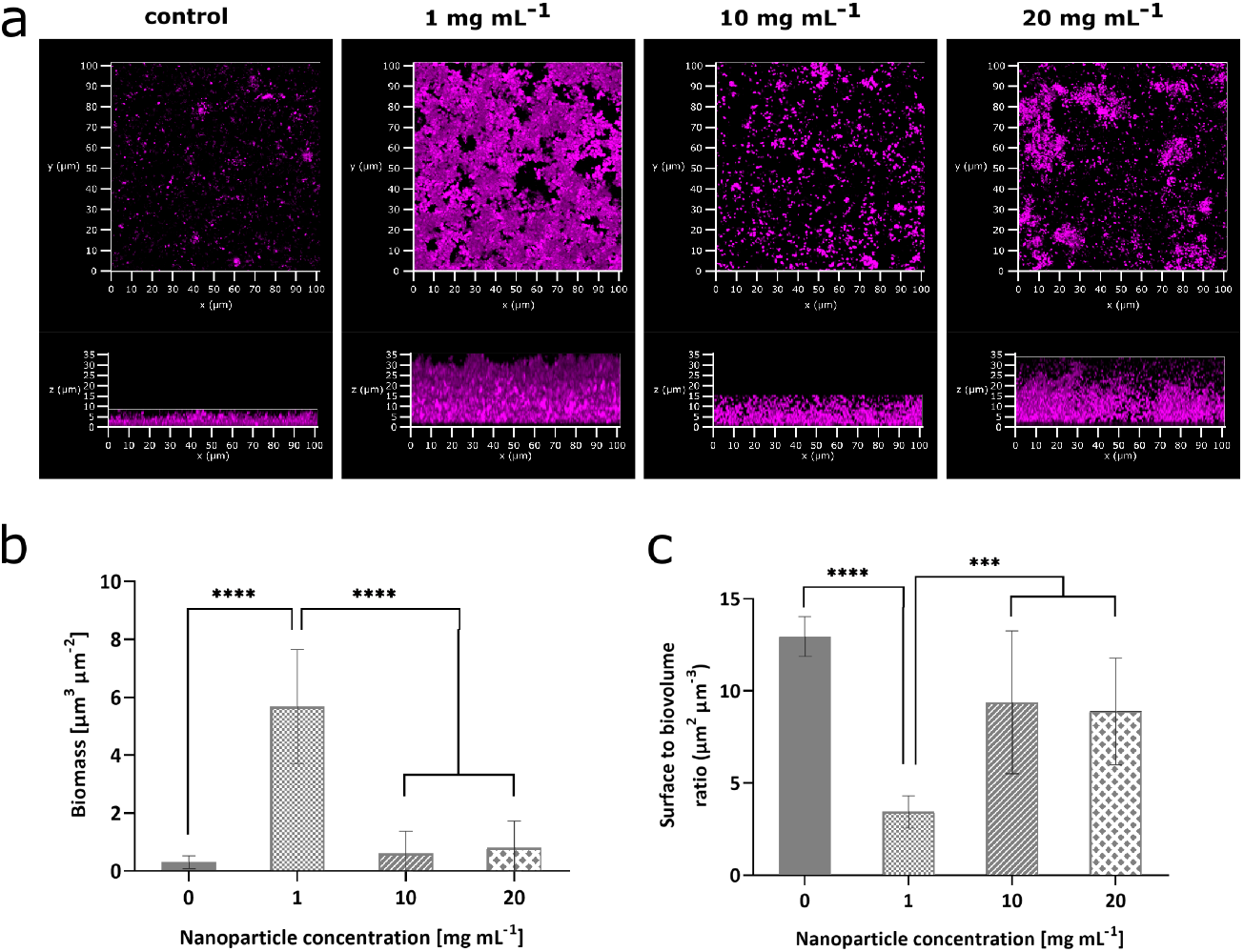
Enhanced *P. aeruginosa* biofilm formation as a result of positively charged nanosensors is concentration dependent. (**a**) Confocal images of *P. aeruginosa* PAO1-N wildtype stained with CellMask™ without the addition of nanoparticles (left panel) or grown with positive polyacrylamide nanosensors at 1 mg mL^1^ (left central panel), 10 mg mL^1^ (right central panel) or 20 mg mL^1^ (right panel). The top row shows a representative 3D view from the top of the biofilm, the bottom row shows a 3D view from the front of the biofilm. Quantitative Comstat analysis of the biofilm images were performed using ImageJ. Graphs represent (**b**) biomass and (**c**) surface to biovolume ratio. Error bars represent standard deviation measured for different biofilm images, where n=10 (control), n=8 (1 mg mL^1^), n=12 (10 mg mL^1^) and n=13 (20 mg mL^1^).

### Polyacrylamide nanosensors rapidly penetrate an established biofilm

To investigate if nanosensors can penetrate an already established *P. aeruginosa* biofilm, nanosensors were added to a 48-h matured biofilm. Penetration of the sensors was assessed using fluorescence CLSM. To ensure that a thick biofilm could be used for the analysis, the bacteria were pre-grown with positively charged nanoparticles without fluorophores. After imaging the biofilm, positively charged or neutral nanosensors were added to the cultures and the biofilms were immediately imaged again without washing. Penetration of the biofilm by the nanosensors was assessed by tracking the fluorescence of the nanosensors throughout the thickness of the biofilm. Fluorescence within the whole biofilm was observed after 3 min following the addition of the nanosensors suggesting that the nanosensors enter the biofilm very quickly and without obvious resistance. This behaviour was observed for both the neutral and the positively charged nanosensors.

Interestingly, when the biofilms were grown initially without the positive nanoparticles, the structure of the biofilm was disrupted when positively charged nanosensors were added but not when neutral nanosensors were added, supporting the observation that growing PAO1-N together with the positive nanoparticles results in a denser biofilm (Fig. 3).

**Fig. 3.**
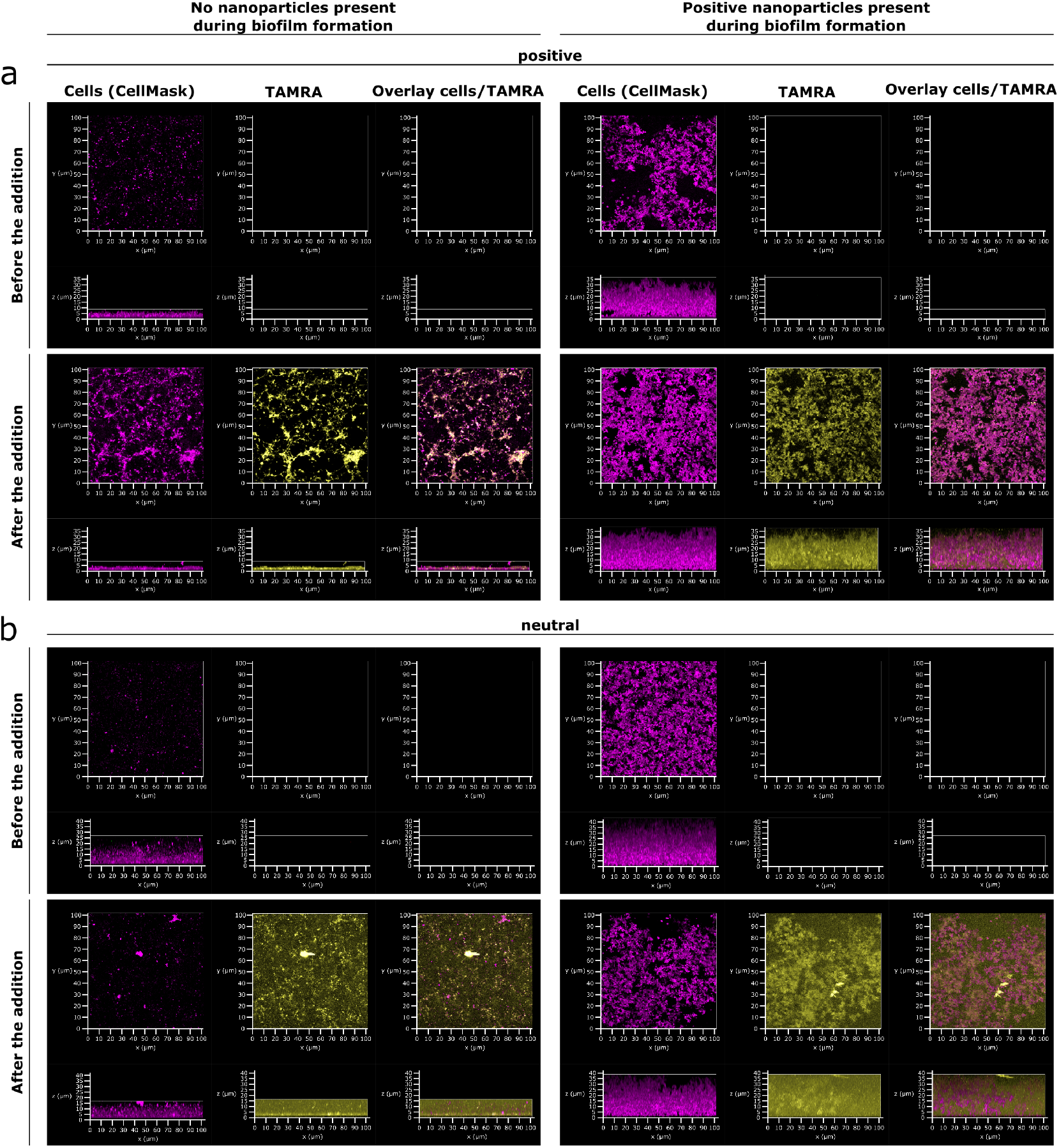
Neutral as well as positive nanosensors can penetrate established *P. aeruginosa* biofilms under static conditions. (**a**) Confocal images of *P. aeruginosa* PAO1-N grown without (left panel) or with (right panel) 1 mg mL^−1^ positive nanoparticles without fluorophores for 48 h. Images show cells stained with CellMask (magenta) and TAMRA fluorescence of nanosensors (yellow). For the purpose of these images TAMRA has been false coloured to yellow to facilitate visualisation of nanoparticle distribution. Top panel = Before the addition of positively charged pH-sensitive polyacrylamide nanosensors. Bottom panel = After the addition of positively charged pH-sensitive polyacrylamide nanosensors to a final concentration of 1 mg mL^−1^. Left panel = (**b**) Confocal images of *P. aeruginosa* PAO1-N with same conditions as in (**a**), but with the addition of 1 mg mL^−1^ neutral pH-sensitive polyacrylamide nanosensors.

In addition to a static biofilm, the flow cell BioFlux system was used to investigate penetration of biofilms by both neutral and positive nanosensors (Fig. 4). After initially growing the biofilm for 12 h in the absence of nanosensors, the inlet of the flow medium was switched to a medium containing nanosensors for 5 h followed by a switch back to medium without nanosensors for another 5 h. CLSM at the end point showed that there were nanosensors incorporated within the biofilm indicating penetration of the biofilm by the nanosensors. Neither neutral nor positive nanosensors changed the biofilm structure or thickness when grown within the BioFlux system.

**Fig. 4.**
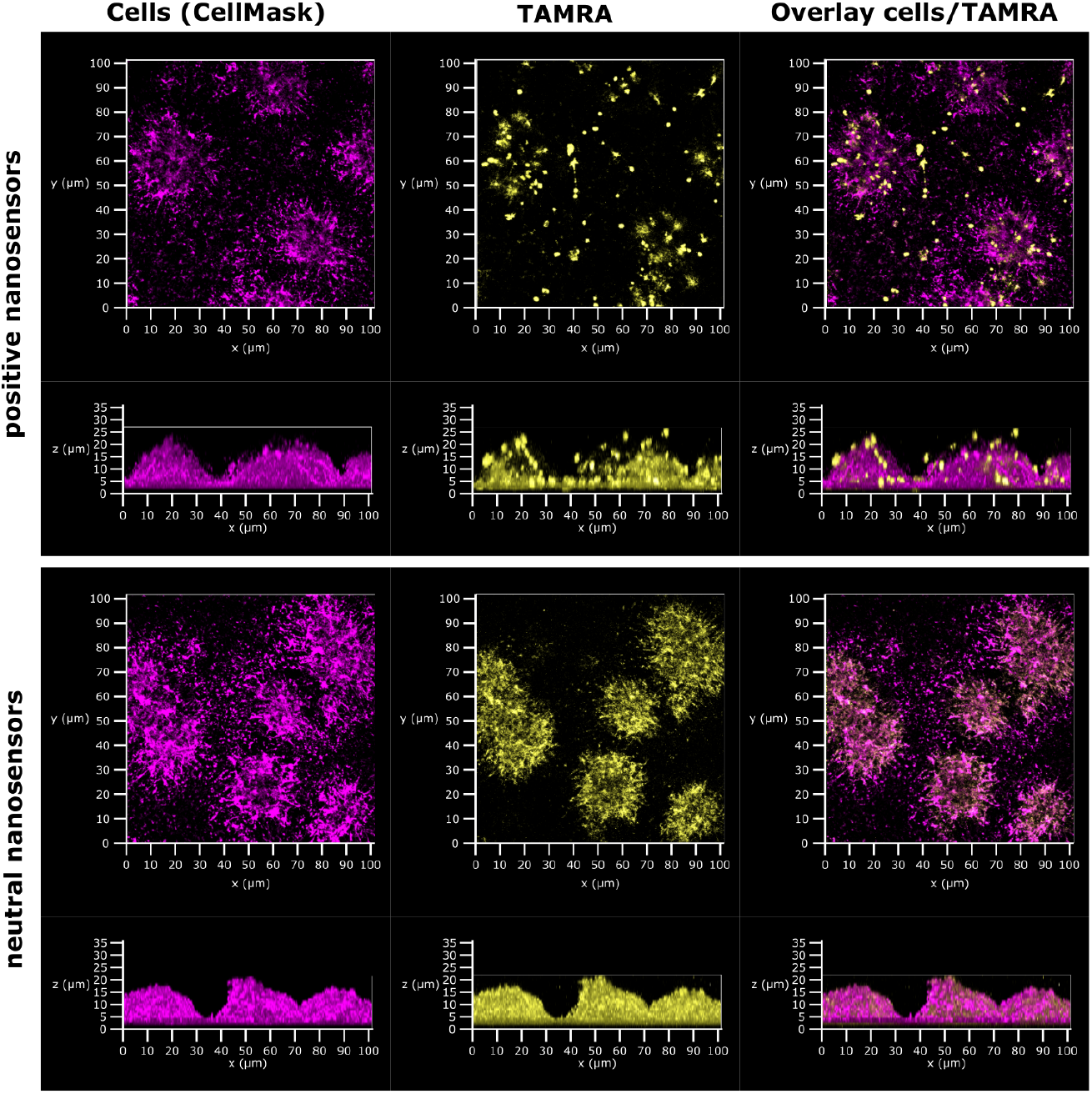
Penetration of established *P. aeruginosa* biofilms by nanosensors under flow conditions was independent of the charge of the nanosensors. *P. aeruginosa* biofilms were initially grown for 12 h without nanosensors before the medium was switched to medium with 5 mg mL^−1^ positive or neutral nanosensors for 5 h followed by a switch back to medium without nanosensors for another 5 h. Confocal images show nanosensor fluorescence (TAMRA, yellow) within the biofilm after the final wash step for both nanosensors (positive and neutral). For the purpose of these images TAMRA has been false coloured to yellow to facilitate visualisation of nanoparticle distribution.

### pH changes can be observed during biofilm formation in a flow cell system at a microcolony level using polyacrylamide nanosensors and time-lapse imaging

Time-lapse imaging together with the nanosensor technology was used to investigate pH changes during biofilm formation of *P. aeruginosa* PAO1-N over time in a BioFlux flow cell system. Images were taken every 15 min for 16 h using a Nikon widefield microscope. A red region of interest indicates a low pH, due to diminishing signal from OG & FAM under acidic conditions, whereas green to orange indicate relatively higher pH values, due to an increase in fluorescence from OG and FAM. During the first couple of hours after microcolony formation starts, the pH increases slightly as observed by a colour shift from a more orange colour (lower pH) towards a more greenish colour (higher pH).

Once the microcolonies started to form after 10 h, the colour around the microcolonies changed to red indicating an acidification during the microcolony formation. Red streaks could be observed after 13 h coming from the microcolonies and being taken away by the flow (Fig. 5 a). Subsequently, the colour continued changing towards red until the whole biofilm was acidified. This observation was confirmed by analysing the fluorescence intensity of the green channel (pH sensitive dyes) and red channel (reference dye) using ImageJ (Fig. 5 b). Over the first hours when microcolonies started to form, the fluorescence intensity ratio of OG/FAM and TAMRA increased slightly from 1.5 to 1.6, indicating a pH increase. The fluorescence intensity around the microcolonies starts to decrease after 10 h before the intensity of the medium declines. After 16 h of biofilm growth, the fluorescence intensity ratio around the microcolonies as well as within the medium reaches 1.25, indicating an acidification of the whole biofilm. A video of the time-lapse imaging can be found in supplementary (Supplementary Movie 1).

**Fig. 5.**
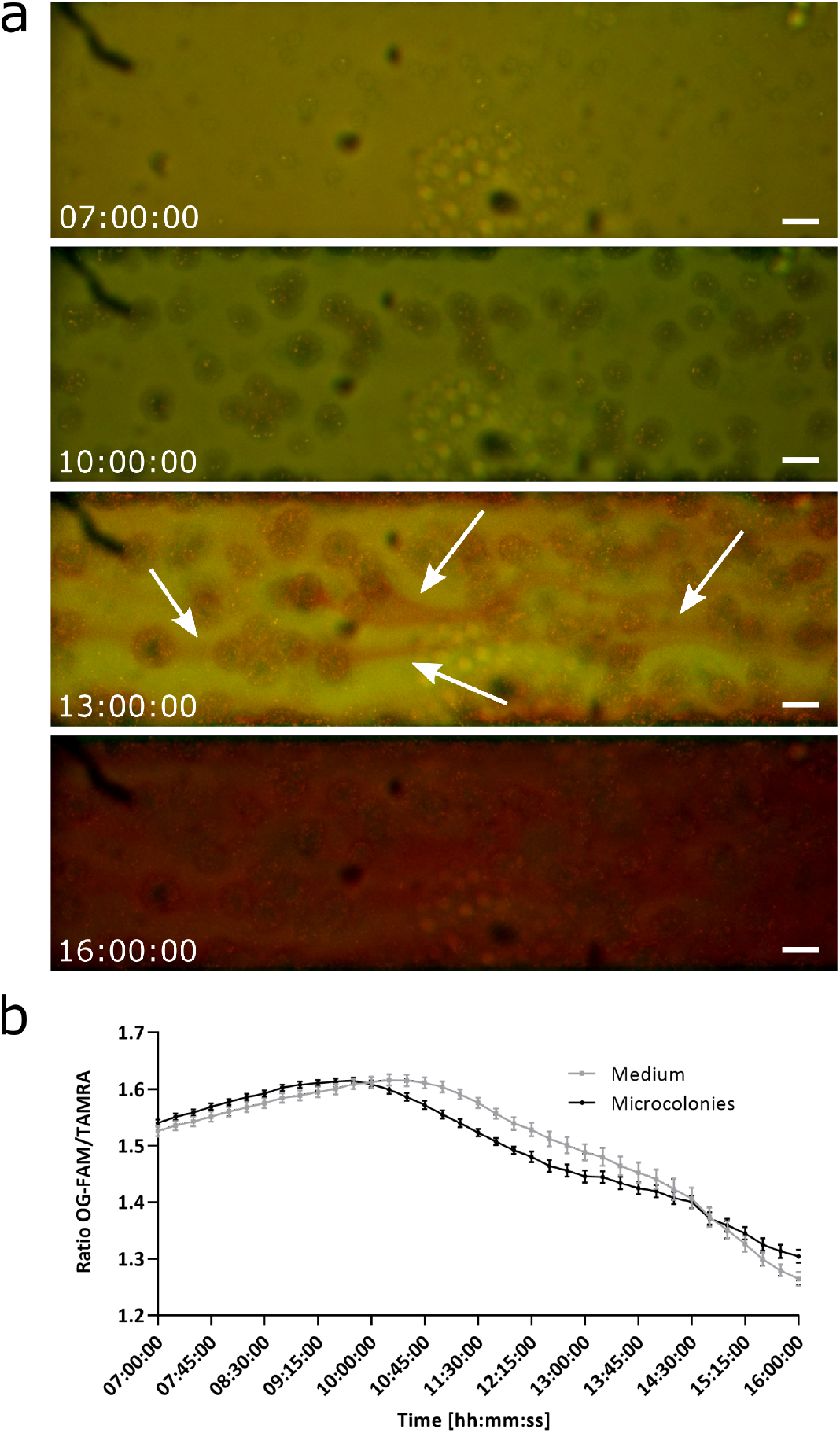
A streamer of acidic pH is evident in the downstream flow of *P. aeruginosa* biofilm microcolonies. Time-lapse images of a *P. aeruginosa* biofilm grown in a BioFlux flow cell and imaged using a fluorescence Nikon widefield microscope. (**a**) Time-lapse imaging of a *P. aeruginosa* biofilm grown in a BioFlux flow cell after 7 h (start of microcolony formation), 10 h, 13 h and 16 h. Red streaks being released downstream by the microcolonies into the medium after 13 h of growth are indicated by the white arrows. Positively charged pH-sensitive polyacrylamide nanosensor were included at 5 mg mL^−1^ to visualize the red streaks. Scale bars represent 50 μm. (**b**) Fluorescence intensity of OG/FAM and TAMRA was measured using wide field images in ImageJ and the ratio is plotted against the time of incubation starting after 7 h when microcolony formation occurred. Error bars represent standard error measured for different areas of the images, where n=6. To view the movie, see Supplementary Movie 1.

### Single microcolonies of a PAO1-N biofilm show pH variation from a more acidic environment within the core to a more neutral pH on the outside surface

CLSM was used to look at the pH variation of single microcolonies of a *P. aeruginosa* PAO1-N biofilm grown within a BioFlux flow cell. The biofilm was imaged in 1 μm steps up to 25 μm. The 3D Z-stack shows a colour change from red within the core of the microcolonies towards yellow at the outer parts (Fig. 6 a). After calibration of the nanosensors using pH buffers ranging from pH 3 to 8, pH maps and a 3D rendering of the Z-stack of the biofilm were generated using the software MatLab ^65^. The values of the pH maps correlate with the observations from the 3D Z-stack showing a low pH within the microcolonies (~pH 3.5 – 4.5) and a more neutral pH towards the outside (pH 6.0 – 7.0) (Fig. 6 b and c). A video of the 3D rendering can be found in supplementary (Supplementary Movie 2).

**Fig. 6.**
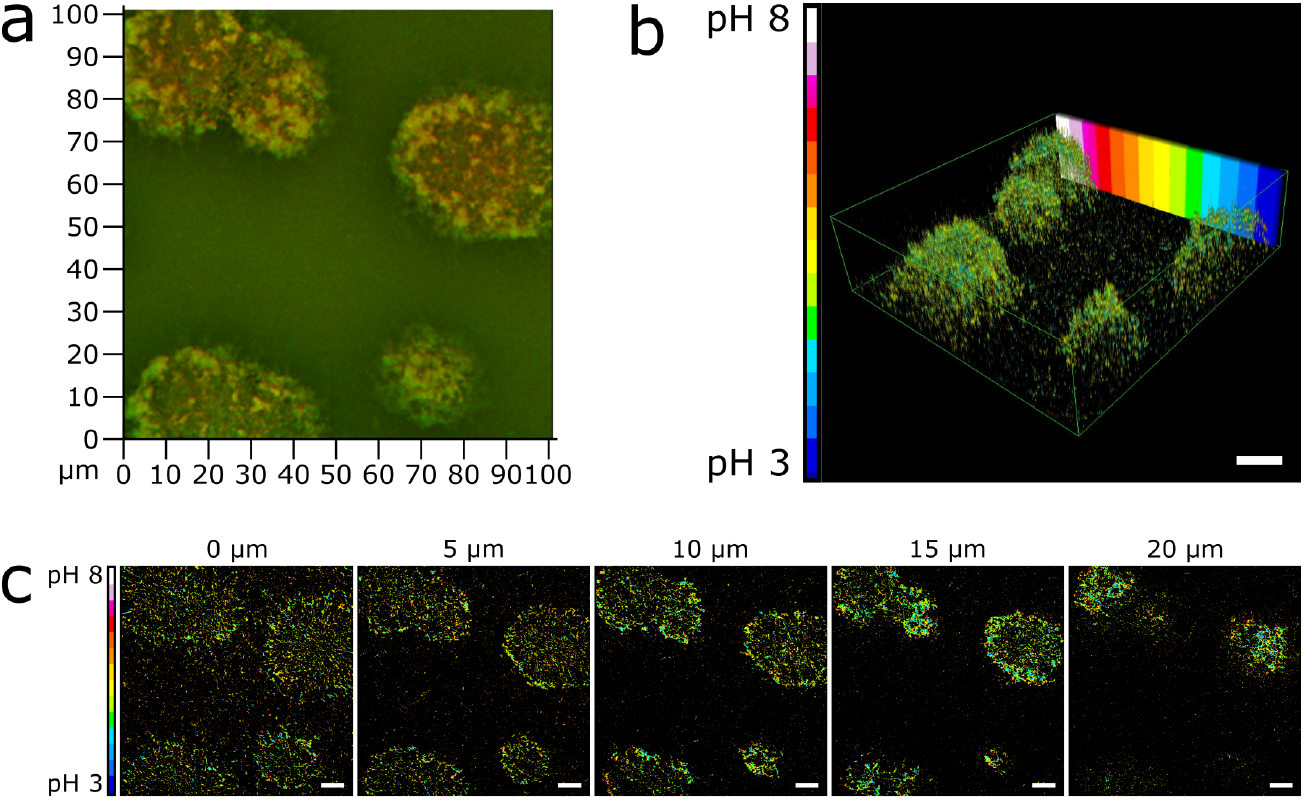
*P. aeruginosa* microcolonies in flow biofilms have an acidic centre. (**a**) The confocal 3D Z-stack image of single microcolony from a *P. aeruginosa* biofilm grown in a BioFlux flow cell system with 1 mg mL^−1^ positively charged pH-sensitive polyacrylamide nanosensors. The pH variation was observed after biofilm growth for 22 h. MatLab analysis of confocal images shows a lower pH of 3.5 – 4.5 within the core of microcolonies and a higher pH of 6.0 – 7.0 at the outer parts. Images represent 3D rendering of the biofilm Z-stack (**b**) and individual pH maps at the bottom of the biofilm (0 μm) and 5 μm, 10 μm, 15 μm and 20 μm from the bottom into the biofilm (**c**). Scale bars represent 10 μm.

### The addition of glucose drastically reduces the pH of the medium when added to starved planktonic *S. mutans*

Having optimised the inclusion of positive pH-sensitive polyacrylamide nanosensors in *P. aeruginosa* biofilms under both static and flow conditions, we applied our novel tool to the analysis of our second bacterial model with the aim to translate our fundamental observations to an applied scenario. If we could demonstrate acidification of the environment by the oral pathogen *S. mutans* linked to the presence of its preferred sugars, our findings would provide a valuable method to assess potential treatments for dental caries. Bearing this in mind, the response of starved planktonic *S. mutans* to the introduction of fermentable carbon sources (glucose and sucrose) versus a non-fermentable carbon source (xylose and xylitol) was analysed, with the aim of detecting any pH changes in the medium as an indirect measurement of the fermentation of a carbon source.

Fluorescence microscopy revealed that the control with saline remained unchanged throughout the experiment. This was also the case following the introduction of the non-fermentable xylose and xylitol, which resulted in no significant change in the pH during the 30 min period. In contrast, during the same time period, the addition of both glucose and sucrose to the starved cells led to a drastic reduction in pH from ~pH 5.3 to ~pH 3.8 and ~pH 5.2 to ~pH 3.9, respectively (Fig. 7).

**Fig. 7.**
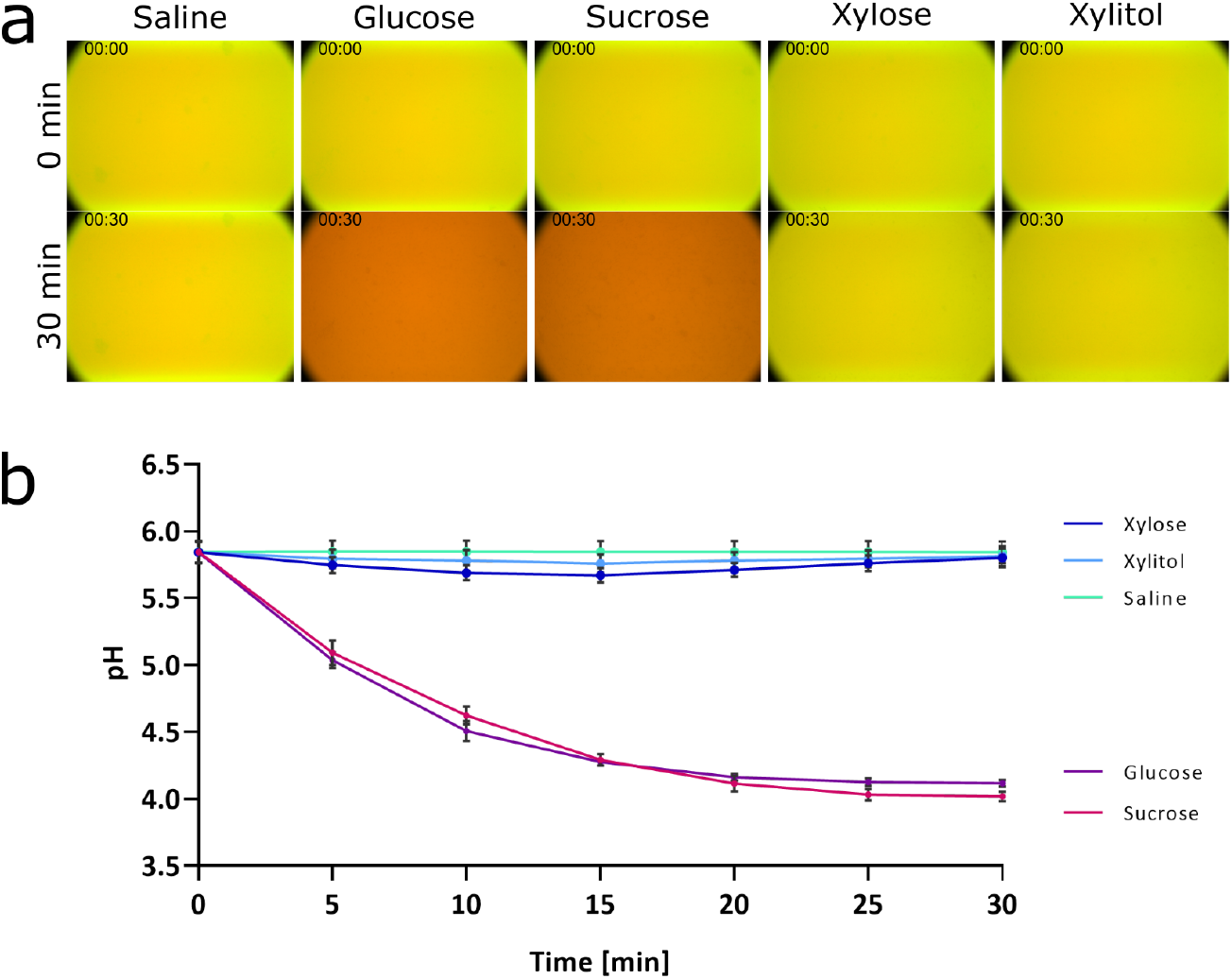
Fermentation of glucose and sucrose, but not xylose, xylitol or saline by *S. mutans* visualised with pH-sensitive nanosensors. Positively charged pH-sensitive nanosensors at a concentration of 1 mg mL^−1^ were added to planktonic *S. mutans* grown overnight and normalized to an OD_600_ of 1. Time-lapse images were taken with a Nikon widefield microscope. (**a**) Representative fluorescence images of cells in saline before the addition of either saline, glucose, sucrose, xylose or xylitol (top panel) to a final concentration of 1% or after 30 minutes (bottom panel). An overlay image of the fluorescence channels for green (OG/FAM) and red (TAMRA) is shown. (**b**) The pH of each condition at the indicated time points was calculated and plotted against the time of addition. Error bars represent standard deviation measured for different images, where n=9.

## Discussion

This study has shown that pH-sensitive polyacrylamide nanosensors penetrate biofilms of *P. aeruginosa*, and report on pH modulation for both our chosen model bacteria, *P. aeruginosa* and *S. mutans.* Once nanosensors have been integrated into biofilms, they are capable of real-time pH quantification in static model systems at high temporal, spatial and measurement resolution that permits determination of (i) a pH gradient across the biofilm thickness, and also (ii) an acidic core to microcolonies. For the first time, these studies also include time-lapse imaging that reveals acidification downstream of microcolonies in a flow biofilm model of *P. aeruginosa*. Moreover, in static models, nanosensors with a positive charge supported the formation of thicker *P. aeruginosa* biofilms. Finally, detection of fermentation of preferred sugars by *S. mutans* has the potential to provide the opportunity to test treatments intended to reduce the progression of dental caries. These studies provide a rationale for, and contribute to the understanding of, the substitution to alternate sugars (xylose and xylitol) for patients at high risk of oral health complications, that could lead to dental caries formation.

The selection of appropriate nanosensors, especially with regards to their charge, is crucial for the analysis of microbial biofilms. The nanosensors chosen for this work are well suited to analyse pH changes over time using time-lapse imaging as well as to look at pH variation within single microcolonies using CLSM. The biofilm thickness promoting effect of positively charged polyacrylamide nanosensors upon static PAO1-N biofilms was independent of the incorporation of fluorophores and conferred an enhanced robustness to the biofilms. Penetration of biofilms by nanoparticles usually occurs by diffusion, with particles interacting with the bacteria and the biofilm matrix components ^66,67,68,69^. Due to the negative surface charge of PAO1-N, it is likely that the positively charged nanosensors are interacting directly with the bacteria. This interaction could facilitate the bacteria to attach to each other and form a denser biofilm as seen here. The nanosensors would in this case function as ‘glue’, sticking the cells together and enable biofilm development. When neutral nanosensors or negatively charged particles are used, this interaction is reduced or even reversed most likely due to the same charge of the bacteria and nanoparticles leading to repulsion instead ^44,59^. The enhanced biofilm formation was not observed when biofilms were grown in the BioFlux flow cell system. Due to the constant flow of medium, it could be postulated that the bacteria are less influenced by the nanosensors and therefore their adherence to each other is not modulated.

The increased thickness of static biofilms was dependent on the concentration of the nanosensors, where higher concentrations of up to 20 mg mL^−1^ did not intensify the effect, but instead led to a biofilm which was less uniform and robust compared to the biofilms incubated with the working concentration of nanosensors (1 mg mL^−1^). Nanosensor concentrations below 20 mg mL^−1^ did not affect the growth of planktonic PAO1-N. It follows that a higher concentration could simply result in flooding of the biofilm with positively charged particles. Too many positively charged nanosensors would be predicted to lead to a repulsion between the nanosensors and possibly other positively charged biofilm components, and consequently less adherence between bacteria, which would in turn reduce the robustness of the biofilm and potentially adversely affect its uniformity.

One negatively charged component of a biofilm that plays an important role for the formation of biofilms is extracellular DNA (eDNA). In a study using 0.2 μm positively and negatively charged fluorescent polystyrene nanoparticles with *Burkholderia cepacia* it was shown that the addition of DNase to the biofilm enhanced the diffusion coefficient of the positively charged nanoparticles, suggesting that there is an interaction between positively charged nanoparticles and the eDNA. This could further increase the ‘glue’ effect and enhance biofilm formation ^70^. Since the polyacrylamide nanosensors used in our study penetrated biofilms so rapidly (within minutes), diffusion gradients would be challenging to measure, and because eDNA deficient mutants of *P. aeruginosa* fail to form biofilms, further investigation of matrix-nanosensor interactions were beyond this study, and will form part of future investigations.

When positively charged nanosensors were added to biofilms that were initially grown without nanosensors, the biofilm growth structure was disrupted. This behaviour was not seen when biofilms were grown with positively charged nanoparticles from the beginning. One hypothesis that could be drawn from this, is that biofilms initially grown with positive nanoparticles were more robust. The less robust biofilms formed in the absence of positively charged nanoparticles might be susceptible to disruption upon introduction of positively charged nanoparticles capable of interacting with the bacterial cell surface or matrix components. This has also been observed by Li *et al.* who used positively charged block copolymer nanoparticles to disperse established biofilms of multidrug-resistant Gram-positive bacteria ^71^. The situation appears to differ in flow cell biofilms since both neutral and positively charged nanosensors were able to penetrate an already established biofilm that had been grown for 12 h. Even after changing the flow cell back to a medium without nanosensors, the nanosensors were still detectable and incorporated within the biofilm. This has the practical application for future investigation as it indicates that the nanosensors can be added at later timepoints for pH analysis if needed.

Being able to increase the thickness of biofilms has beneficial implications. For example, the positively charged nanosensors could be employed in future studies to elucidate the mechanistic steps of biofilm formation by combining them with mutants defective in defined aspects of biofilm formation. On the other hand, neutral nanosensors could be the better choice to study biofilms that closely mimic the natural physiological or environmental conditions, as no structural change in biofilm formation behaviour was observed when they were applied to our models.

Extrapolating from the influence of nanoparticle charge on biofilm formation, it is interesting to speculate that the charge of antimicrobials could influence the structure of biofilms. For example, antimicrobials with a positive charge could not only have biocidal effects such as benzalkonium chloride ^72^, but also interact with cells in a way that could enhance biofilm formation or dispersal. It would follow that this might affect medical treatment or anti-biofouling regimens. It will therefore be interesting to monitor local pH modulation during antimicrobial application to developing or established biofilms and correlate this with biofilm architectural changes.

This study has shown for the first time, that pH-sensitive polyacrylamide nanosensors can be used to visualise pH changes over time during biofilm formation of PAO1-N within a BioFlux flow cell system at a microcolony level. At the beginning of the biofilm formation, the pH increases very slightly as the bacteria start to adapt to the environment. After 10 h, acidification of the biofilm was observed during microcolony formation and subsequently while the biofilm continued to form. The acidic streaks that were released by the microcolonies could be exoproducts from the bacteria, produced during metabolism ^44^. Another possibility is that acidic molecules are released by the cells to help form the biofilms. These molecules could for example be eDNA or exopolysaccharides (EPS). It could also be that the cells are actively lowering the pH when biofilm formation is initiated, to facilitate the process, for example to enable better attachment or to increase the turn-over and release of quorum sensing molecules like AHLs ^35,73^.

In addition to the pH changes observed over time within the whole biofilm, pH variation was also detected within single microcolonies, showing a more acidic environment within the core of the microcolonies and a more neutral pH at the outside edge of the colonies. The acidic cores could result from the production and release of acidic metabolites during growth and biofilm formation. Furthermore, oxygen will likely be depleted within the core and no fresh oxygen would reach the inner part of the microcolonies resulting in fermentation processes of the cells, which would then lead to the production of acidic products, reducing the pH ^26^.

To provide a translational benefit of the pH-sensitive nanosensors, they were dispersed within the medium of starved planktonic *S. mutans*, to measure changes in fluorescence intensity and calculate the resultant external pH changes. By utilising the well-studied biological system of *S. mutans*, where carbohydrate supplementation can induce a pH response, the effectiveness of the pH sensitive polyacrylamide nanosensors as a real-time tool to accurately determine pH changes in extracellular media was illustrated. Oral bacteria including *S. mutans*, must adapt to the shifting availability of carbohydrates caused by changes in host behaviours. These can be brought on through the fluctuation of diets, as well as through host secretions of glycoproteins and those carbohydrates produced by the oral microbiome itself ^74^. *S. mutans* is therefore versatile in the carbohydrates it can utilise ^75^. With the addition of simple mono- and disaccharides such as glucose or sucrose, a reduction in the pH was observed, as *S. mutans* is capable of metabolising both glucose and sucrose via glycolysis to produce pyruvate ^16^. The resultant pyruvate is metabolised to form L-lactate which would be secreted in the form of lactic acid, leading to the reduction of the pH of the medium ^76^. However, with the addition of xylose or its derivative, xylitol, the external pH was unchanged throughout the experiment. This was a result of both xylose and xylitol being non-fermentative by *S. mutans* ^77^. Xylose is initially taken up and reduced to xylitol, which is then phosphorylated to xylitol-5-phosphate (X5P). *S. mutans* is unable to metabolise X5P further, and as a result X5P is accumulated intracellularly ^78^. This accumulation of X5P has been attributed to the inhibition of glycolytic enzymes; leading to the repression of acid production ^78,79^. However, Takahashi and Washio ^80^ were able to show that the presence of X5P had no effect on acid production when supragingival plaques were rinsed with glucose after an initial application of xylitol. This would imply that xylitol is simply a non-fermentative sugar alcohol rather than an inhibitor.

A practical application of our nanosensor system could be envisioned for testing oral hygiene products or sweetener alternatives in soft drink production, in order to determine the impact on pH production. The nanosensors would provide flexibility, as a screening tool to measure pH changes in high throughput fluorescence assays. Alternatively, the nanosensors could be used as a more focused tool, to track changes in the pH microenvironment of established oral biofilms over time, to map 3D pH microenvironments as highlighted by our analysis in the flow cell.

Future work will also explore the molecular mechanisms underpinning the detected acidification of the biofilms and microcolony centres. For example, different fluorophores could be used to stain eDNA or EPS components to investigate whether either correlates with the acidic streaks observed. Furthermore, adding or changing dyes in the nanosensors could enable more powerful tools to be developed for monitoring broader environmental microniche changes within biofilms in real-time and at high resolution.

## Materials and Methods

### Materials

Oregon Green^®^ 488 carboxylic acid (OG), 5-(and-6)-carboxyfluorescein (FAM) and 5-(and-6)-carboxytetramethylrhodamine (TAMRA) were obtained from Invitrogen™, USA. Acrylamide (≥99%), N,N′-methylenebis(acrylamide) (bisacrylamide, 99%), dioctylsulfosuccinate sodium (AOT), ammonium persulphate (APS, ≥98%), 3-acrylamidopropyltrimethyl ammonium hydrochloride (ACTA, 75 wt. % in H_2_O), polyoxyethylene(4)lauryl ether (Brij L4^®^), sodium tetraborate decahydrate (≥99.5%) and N,N,N,N-tetramethyl-ethylenediamine (TEMED, 99%) were purchased from Sigma Aldrich, US. N-(3-Aminopropyl) methacrylamide hydrochloride (APMA, >98%) was obtained from Polysciences Inc, Germany. Hexane, ethanol absolute (99.5%) and phosphate buffer saline (PBS) were obtained from Fisher Scientific, UK. Unless otherwise mentioned, all the chemicals that used throughout this study were of analytical grade.

### Minimal medium (M9)

Minimal medium (M9) was prepared by using 200 mL of 5x M9 salts (Sigma Aldrich), 2 mL MgSO_4_ (1M), 100 μL CaCl_2_ (1M) and 40 mL Succinate (0.5 M) in a total of 1 L.

### Bacterial strains and growth culture

The bacterial strains used in this study were PAO1-N wildtype ^81^ and *Streptococcus mutans* NCTC 10449 ^82^. Lysogeny Broth (LB) ^83^ was used as standard growth medium (liquid or solid as agar plate) for PAO1 and Brain Heart Infusion (BHI) for NCTC 10449. *P. aeruginosa* strains were grown at 37°C for 16-20 h, *S. mutans* strains were grown at 37°C for 48 h. Liquid cultures were prepared in 5 mL liquid media, inoculated with a single bacterial colony and shaken at 200 rpm (PAO1) or incubated in a static incubator (NCTC 10449) overnight. Strains were stored for the longer term as glycerol cultures which were made by using 500 μL overnight culture with 500 μL sterile 50% glycerol and stored at 80°C.

### Static biofilm growth of PAO1

Overnight cultures of *P. aeruginosa* grown in LB were used to spin down 1 mL of cells in a fresh Eppendorf tube at 1000 x *g* for 1 min. The pellet was re-suspended in 1 mL Phosphate buffer saline (PBS), spun down again at 1000 x *g* and re-suspended in 1 mL of minimal medium (M9 succinate). The OD_600_ was taken and the cells normalised to 0.5 OD_600_ in 1 mL of selected medium. Nanosensors were suspended at 1.5 mg mL^−1^ in selected medium. Polyacrylamide nanosensor suspensions were filter sterilised using 0.22 μm PES filters and 500 μL of medium with nanosensors were combined with 175 μL medium and 75 μL normalised cells to obtain a final concentration of 1 mg mL^−1^ nanosensors and a starting OD_600_ of 0.05. For approaches without nanosensors, another 175 μL medium was added instead of medium with nanosensors. To each well of the 8-well chamber (Ibidi, glass bottom), 300 μL of the suspension was added before the chamber was placed in a box wrapped in aluminium foil and placed in a static incubator at 37°C. After an incubation of 48 h, the media was carefully removed and replaced by fresh medium containing 2.5 μg mL^−1^ CellMask™ Deep Red plasma membrane stain (ThermoFisher Scientific). The biofilms were imaged using a Zeiss confocal laser scanning microscope and the appropriate excitation settings for the fluorescence channels (DAPI = 405 nm, OG/FAM = 488 nm, TAMRA = 555 nm).

### Flow cell biofilm of PAO1 growth using BioFlux

Overnight cultures of *P. aeruginosa* grown in LB were used to set up liquid cultures in 5 mL LB using 100 μL of the overnight culture. The cultures were grown to 0.4-0.8 OD_600_ and normalised to an 0.05 OD_600_ in 1 mL of minimal medium (M9 succinate). Nanosensors were suspended at 5 mg mL^1^ in selected medium and filter sterilised using 0.22 μm PES filters. The BioFlux flow cell (BioFlux 200 48-well low shear plate, 0-20 dynes cm^−2^) was prepared according to the manufacturer’s instructions (Fluxion Biosciences, BioFlux System). For the seeding of the cells, 50 μL of the normalised cells were used with a seeding time of 45 to 60 mins. After the seeding, 1 mL of the prepared medium with or without nanosensors was used as flow medium. The flow was set to 0.25 dyn cm^−2^ and the system was run for 20-24 h. Time-lapse imaging was applied using a Nikon widefield microscope with brightfield and fluorescence channels (OG/FAM excitation = 460 nm, TAMRA excitation = 550 nm). Images were taken every 15 min for 16 h. ^84^

### Sugar challenge with planktonic NCTC 10449 cells

An overnight culture of NCTC 10449 in BHI was spun down and re-suspended in PBS for 30 min as a starvation step. The cells were spun down again and re-suspended in saline before being mixed with positively charged nanosensors in saline to obtain an OD_600_ of 1 and a nanosensor concentration of 1 mg mL^−1^. The suspension was aliquoted into a *Greiner* 24-well, PS, flat glass bottom, black walled microplate using 400 μm per well. Imaging was performed using wide field fluorescence microscopy with the 40×/0.6 lens (40x objective with 1.5x additional magnification). An image was taken (brightfield, OG/FAM excitation = 460 nm, TAMRA excitation = 550 nm) for time point 0 min before 20 μL of either glucose, sucrose, xylose, xylitol (all 1% final conc.) or saline was added to their respective wells. Images were taken every 5 min for 30 min. Additional controls with glucose, sucrose, xylose, xylitol and saline solution added to 1 mg mL^−1^ positively charged nanosensors in saline were also imaged to confirm the absence of any pH change brought on by the solutions themselves.

The calibration was performed using a Greiner 24-well microplate where pH buffers from pH 8 to pH 2.5 were mixed with positively charged polyacrylamide nanosensors for a final concentration of 1 mg mL^−1^. Images were taken from each pH using the same exposure settings as the experiment. The fluorescence intensity ratio from each pH was plotted and the equation of the line calculated to determine the pH values from the fluorescence intensities generated during the experiment.

### Nanosensor preparation

#### Conjugation of the three fluorophores OG, FAM and TAMRA to APMA

A 50 mM (pH 9.5) Sodium tetraborate decahydrate buffer in water was prepared and used to make a 0.002 mmol APMA solution. For each fluorophore, 1 mg was weighed out in separate vials and suspended in 200 μL of the APMA solution. The mixtures were sonicated for 5 min before being incubated on an oscillator and room temperature for 24 h in the dark. The dyes were stored at −20°C for further use.

#### Synthesis of polyacrylamide pH-sensitive fluorescent nanosensors

For the synthesis of the nanosensors, 1.59 g AOT and 3.08 g Brij L4 were weighed out, mixed in a 250 mL round bottom flask, and deoxygenated for 15-20 min using argon while being stirred. A 500 mL round-bottom flask was used to deoxygenate 100 mL hexane for 30 min using argon before 42 mL of the deoxygenated hexane was added to the AOT and Brij solution. The flask was sealed with a stopper and a balloon under an argon atmosphere with continuous stirring. For the synthesis of neutral polyacrylamide nanosensors, 540 mg acrylamide and 160 mg bisacrylamide were dissolved in 1.5 mL deionised water, whereas 513 mg acrylamide, 152 mg bisacrylamide and 119 μL ACTA were used the preparation of positively charged polyacrylamide nanosensors. The APMA-fluorophore conjugates were added to the acrylamide solution using 15 μL OG-APMA, 15 μL FAM-APMA and 60 μL TAMRA-APMA. The mixture was deoxygenated and added to the flask containing the surfactants and hexane using a syringe. After 10 min, 30 μL of APS (10% w/v) and 15 μL TEMED were added. The flask was deoxygenated again, sealed, wrapped in aluminium foil, and incubated on the stirrer for 2 h. After the incubation, the hexane was removed using a rotary evaporator at 30°C and 30 mL of ethanol (100%) was added. The mixture was transferred to a 50 mL falcon tube and spun down at 3800 x *g* for 3 min. The supernatant was discarded, and the pellet re-suspended in 30 mL of ethanol (90% in water). The mixture was spun down again at 3800 x *g* for 3 min and washed twice in 30 mL ethanol (100%) and then suspended in 10 mL of ethanol (100%). The mixture was transferred into a clean 250 mL flask and the ethanol was removed using the rotary evaporator at 30°C before the nanosensors were collected and stored at −20°C in the dark until further use.

### Nanosensor calibration and pH calculation using Matlab

Buffer solutions with a pH ranging from 3.0 to 8.0 in 1.0 steps were prepared using 0.2 M Sodium phosphate (Na_2_HPO_4_) and 0.1 M Citric acid (C_6_H_8_O_7_) as indicated in Supplementary Table 1. The pH was adjusted using 0.2 M Sodium phosphate and 0.1 M Citric acid.

For the pH calibration of the nanosensors in a BioFlux experiment, 10 mg mL^−1^ polyacrylamide were prepared in deionised water and mixed 1:1 with each of the pH buffers to get a concentration of 5 mg mL^−1^. To enable the calculation of a calibration curve, 500 μL of each solution were added to a BioFlux plate and was run for 2 min with a flow rate of 0.25 dyn cm^−2^. Images were taken for each buffer sample using a Zeiss CLSM (OG/FAM = 488 nm, TAMRA = 555 nm). The software MatLab was used to calculate the calibration curve and subsequently to determine the pH of the actual biofilm images taken with the Zeiss CLSM.

### Zeta potential and size measurement of nanosensors

For the determination of the zeta potential, 1 mg mL^−1^ nanosensors were prepared in PBS (10%) and transferred into a disposable folded capillary cell using a 2 mL syringe. For the size determination of polyacrylamide nanosensors, 1 mg mL^−1^ nanosensors were prepared in PBS (10%) and transferred into a cell disposable cuvette. Both the zeta potential and the size were measured using the Zetasizer.

### Widefield microscopy

Time-lapse imaging and planktonic sugar challenge were performed using a Nikon eclipse Ti2-U widefield microscope fitted with a Nikon S Plan Fluor ELWD 20x/0.45, Plan Fluor 10x/0.30 or CFI60 40x/0.6 objective lens. Images were captured by using a CoolSNAP™ MYO CCD Camera connected to Nikon software and analysed using ImageJ.

### Confocal laser scanning microscopy

To generate more detailed images, a Zeiss LSM 700 compact confocal laser scanning microscope was used fitted with HAL 100C lamp for light illumination and a Zeiss alpha-Plan-Apochromat 63x/1.46 Oil objective lens. Images were captured by using AxioCam digital microscope camera connected to ZEN software and analysed using Zen blue software.

## Acknowledgements

We thank Katherine Thompson at Unilever R&D Port Sunlight, Merseyside, UK for providing us with scientific insights to assist our research. This work was partly funded by Unilever plc. This work was also supported by funding from the Biotechnology and Biological Sciences Research Council (BBSRC; Award Number BB/R012415/1). B.B. was a recipient of University of Nottingham PhD scholarship shared jointly between the Schools of Life Sciences and Pharmacy, and M.P. was funded by BBSRC iCASE PhD studentship through grant BB/M008770/1. This work was supported by a Nottingham Research Fellowship from the University of Nottingham, UK (V.M.C). We also thank Dean Walsh for useful discussions and help with the nanosensor preparation and James Brown for providing strains.

## Author information

### Contributions

The study was designed by B.B., M.P., J.W.A. and K.R.H. with contributions from V.M.C. B.B. performed all experiments involving *Pseudomonas aeruginosa* and M.P. carried out experiments on *Streptococcus mutans*. B.B. and M.P. performed the data analysis. B.B., M.P., J.W.A. and K.R.H. wrote the manuscript and V.M.C. was involved in critically reviewing the paper.

