## Supplementary Table 1 and Supplementary Figure 1 for "Fluorescent Nanosensors Reveal Dynamic pH Gradients During Biofilm Formation"

**Supplementary Figure 1. No growth inhibition of nanosensors concentrations below 25 mg mL$^{-1}$.**

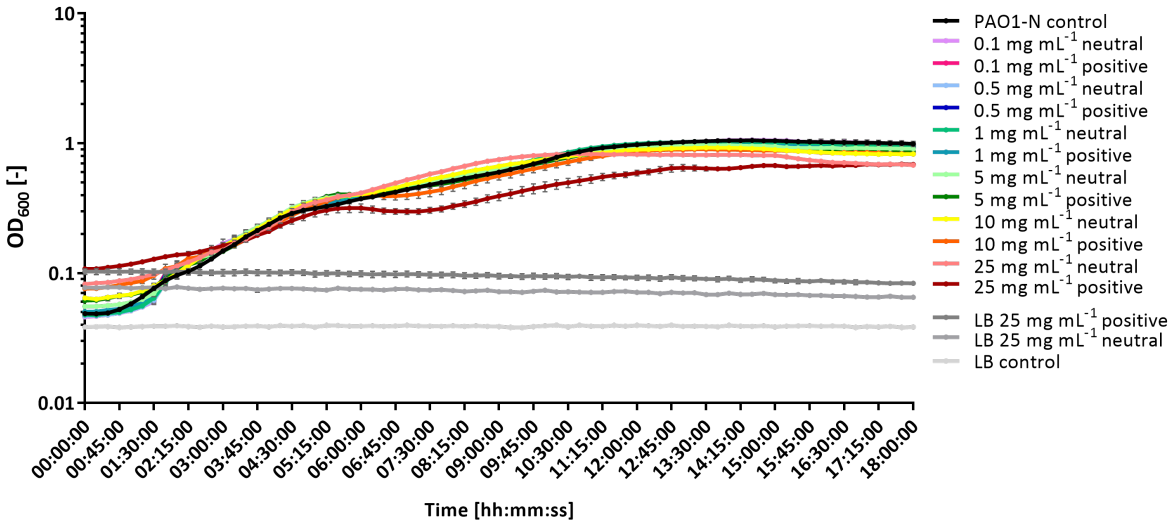


**Supplementary Table 1. pH buffer pH buffer preparation using 0.2 M Sodium phosphate ($Na_2HPO_4$) and 0.1 M Citric acid ($C_6H_8O_7$).**

| pH | $Na_2HPO_4$ [mL] | $C_6H_8O_7$ [mL] | $H_2O$ [mL] |
|---|---|---|---|
| 3.0 | 2.04 | 7.96 | 10 |
| 4.0 | 3.86 | 6.14 | 10 |
| 5.0 | 5.14 | 4.86 | 10 |
| 6.0 | 6.42 | 3.58 | 10 |
| 7.0 | 8.72 | 1.3 | 9.98 |
| 8.0 | 9.765 | 0.24 | 9.99 |

*$Na_2HPO_4$* 0.2 M Sodium phosphate, *$C_6H_8O_7$* 0.1 M Citric acid, $H_2O$ water
